# Lipid interactions of an actinoporin pore-forming oligomer

**DOI:** 10.1101/2020.10.02.323956

**Authors:** A. Sepehri, B. Nepal, T. Lazaridis

**Affiliations:** City College of New York; City College of CUNY

**Keywords:** Actinoporins, fragaceatoxin C, toxins, sphingomyelin, molecular dynamics

## Abstract

The actinoporins are cytolytic toxins produced by sea anemones. Upon encountering a membrane, preferably containing sphingomyelin, they oligomerize and insert their N-terminal helix into the membrane, forming a pore. Whether sphingomyelin is specifically recognized by the protein or simply induces phase coexistence in the membrane has been debated. Here, we perform multimicrosecond molecular dynamics simulations of an octamer of fragaceatoxin C, a member of the actinoporin family, in lipid bilayers containing either pure 1,2-Dioleoyl-sn-Glycero-3-Phosphocholine (DOPC) or a 1:1 mixture of DOPC and palmitoyl sphingomyelin (PSM). The complex is highly stable in both environments, with only slight fraying of the inserted helices near their N-termini. Analyzing the structural parameters of the mixed membrane in the course of the simulation we see signs of a phase transition for PSM in the inner leaflet of the bilayer. In both leaflets, cross-interactions between lipids of different type decrease over time. Surprisingly, the aromatic loop thought to be responsible for sphingomyelin recognition interacts more with DOPC than PSM by the end of the simulation. These results support the notion that the key membrane property that actinoporins recognize is lipid phase coexistence.

**SIGNIFICANCE STATEMENT:** The mechanism of selectivity of naturally produced toxins for their target membranes is not well understood. For example, actinoporins, toxins produced by sea anemones, have been reported to selectively target sphingomyelin-containing membranes. Whether they bind this lipid preferentially or recognize the phase coexistence that sphingomyelin induces is not clear. This work examines this issue by long computer simulations of an actinoporin oligomer embedded in lipid bilayers and finds no preferential interactions of the protein with sphingomyelin. Instead, the simulations show signs of phase separation, suggesting that phase coexistence is the key property that actinoporins recognize.

## INTRODUCTION

The actinoporins are a family of ~20 kDa cytolytic toxins produced by sea anemones (1, 2). The best-studied members of the family are equinatoxin II, sticholysin I and II, and fragaceatoxin C. These toxins are produced as soluble monomers, whose structure is a β-sandwich flanked by two helices (3–7). Upon encountering a target membrane they rearrange into a pore-forming oligomer. The rearrangement involves primarily the conserved amphipathic N-terminal (Nt) helix which extends and inserts into the membrane (8, 9). Early evidence pointed toward a trimeric or tetrameric oligomer forming toroidal pores (5, 10, 11), but more recent crystal structures of fragaceatoxin C in lipidic environments showed a nonameric prepore (12) and an octameric pore with the Nt helices forming a funnel-shaped pore (7). The Nt helices by themselves have some pore-forming activity that is similar to that of the entire toxin in the case of sticholysin (13, 14), but lower in the case of equinatoxin II (15).

The fact that preincubation of actinoporins with sphingomyelin (SM) inhibits them, led to the proposal that SM is their receptor in the target membranes (16, 17). SM recognition was attributed to aromatic residues in the membrane-facing β7-β8 loop, based on measurements of lipid binding in solution, thermal stability, SPR to liposomes, and mutations of aromatic residues (18). However, a molecular understanding of SM recognition is lacking. SM has the same headgroup as PC, so any recognition of one over the other must take place either beneath the headgroup or via an indirect effect of the lipid backbone on the disposition of the headgroup. Other groups found that SM is preferred but not required for ion channel formation (19). More recent articles reported that actinoporins also function on PC/cholesterol membranes that exhibit phase coexistence (20, 21). This conclusion was corroborated by work on GUVs, which showed that phase coexistence is necessary for pore formation (22). Studies with the Droplet Interface Bilayer technique found that the pores occur in the Ld phase, after first binding at the domain boundaries (23), although the GUV study found that the pores prefer the Lo phase (22).

Computer simulations could provide important insights into the mechanism of lipid recognition (24). However, to our knowledge, the issue of SM recognition has not yet been addressed by simulation methods. Recent work on a tetramer of the Nt helices of sticholysin I and II was conducted in POPC/SM & PC/SM/PE membranes (25) and found enrichment of the pore region in PE. SM did not appear to play a role in Nt helix pore formation. Another set of simulations was done on full equinatoxin and the 1-32 peptide on the surface of DMPC or DMPC/SM bilayers (26). Some preference of the Ct of the peptide for SM was noted. Molecular dynamics (MD) simulations of equinatoxin II interacting with DPC and SM micelles have also been reported (27). Binding of single lipids to sticholysin II in solution was evaluated using MM-GBSA (28).

To gain insights into the role of SM in actinoporin function, we conducted multimicrosecond MD simulations of the entire Fragaceatoxin C (FragC) octamer (7) in DOPC and in a 1:1 PSM/DOPC mixture (Fig. 1). Suprisingly, we find that the aromatic loop thought to be responsible for SM recognition interacts preferentially with DOPC rather than PSM. In contrast, in the inner leaflet, the N-terminus interacts preferentially with PSM. We also see evidence of incipient phase separation in the lipid mixture. The results are discussed in the light of available experimental data.

**Fig. 1.**
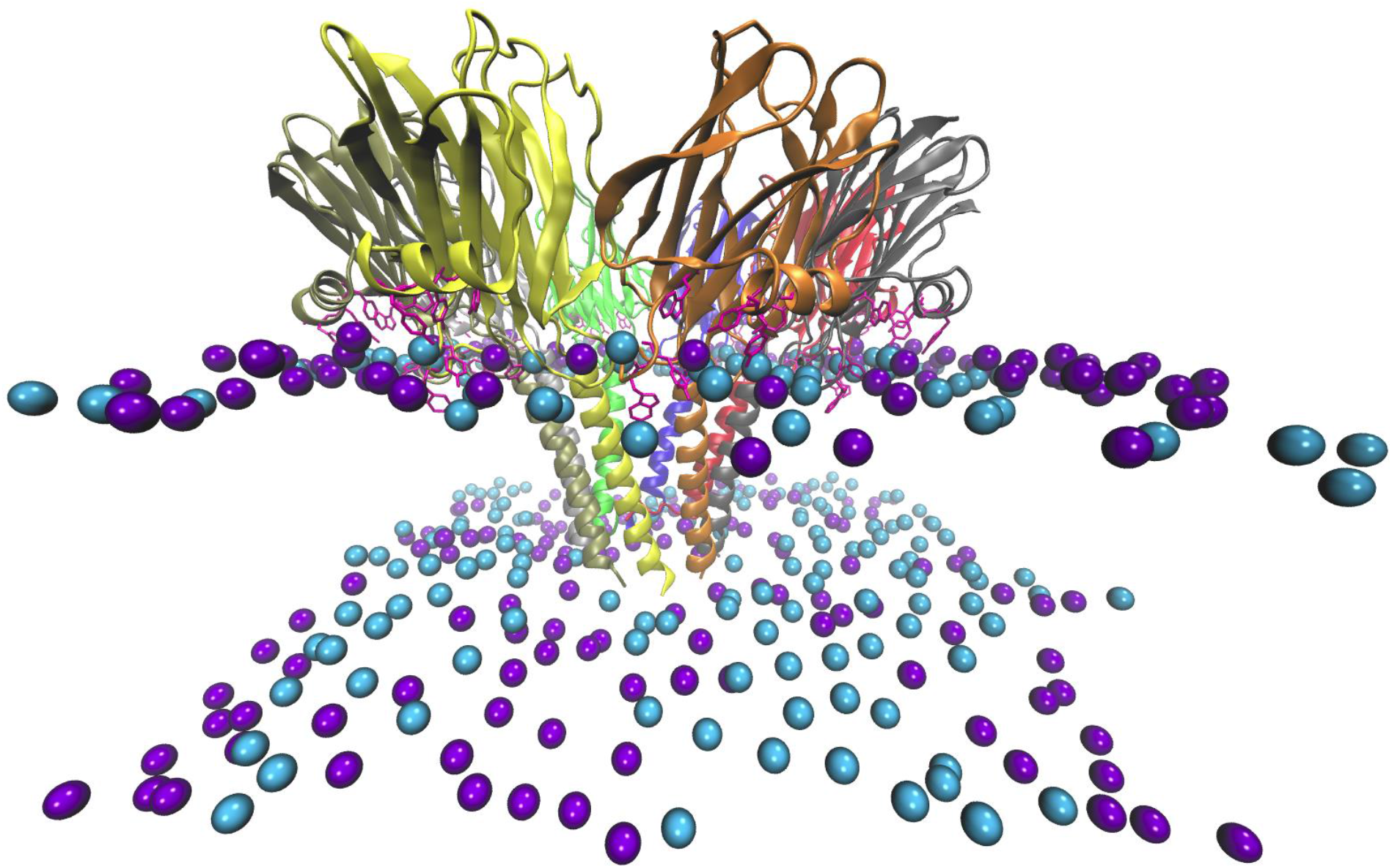
The fragaceatoxin octamer in the mixed bilayer. The N-terminus is on the surface of the inner (lower) leaflet and the aromatic loop on the surface of the outer (upper) leaflet. Cyan and purple spheres are DOPC and PSM headgroups, respectively.

## METHODS

The starting structure of all simulations was the FragC octamer in lipidic mesophase (PDB ID 4TSY). In this PDB file the three first residues, Ser1, Ala2 and Asp3, are missing, probably due to disorder. These residues were built in an extended conformation. The eight protomers are named A through H and presented with these colors in figures: A (dark blue), B (red), C (black), D (orange), E (yellow), F (olive), G (grey), and H (bright green). Two membrane systems were considered: pure 1,2-dioleoyl-sn-glycero-3-phosphocholine (DOPC) and an equimolar mixture of DOPC and palmitoylsphingomyelin (PSM).

The CHARMM-GUI server (29) was used to build initial configurations. The lipid bilayer was built around the eight N-terminal helices. Because of the funnel shape of the pore, the number of lipids in the leaflet close to the peptide’s N-terminus (inner leaflet) is higher than that in the other (outer) leaflet. Then, a 17.5 Å water slab and balancing counter ions were added to the system. Table 1 gives the basic statistics of the two systems. The MD simulations were run on the Anton 2 special-purpose computer (30) using the CHARMM 36 force field (31, 32), TIP3P water (33) and multigrator method (34). Simulations were run in the NPT ensemble at 310 K and 1 bar using the Nose-Hoover thermostat (35) and the MTK barostat (36). Anton implements bond constraints (37) that allow solving the equations of motion numerically with a 2.5 fs timestep. The cutoff distances for nonbonded interactions were determined automatically by Anton.

**Table 1.**
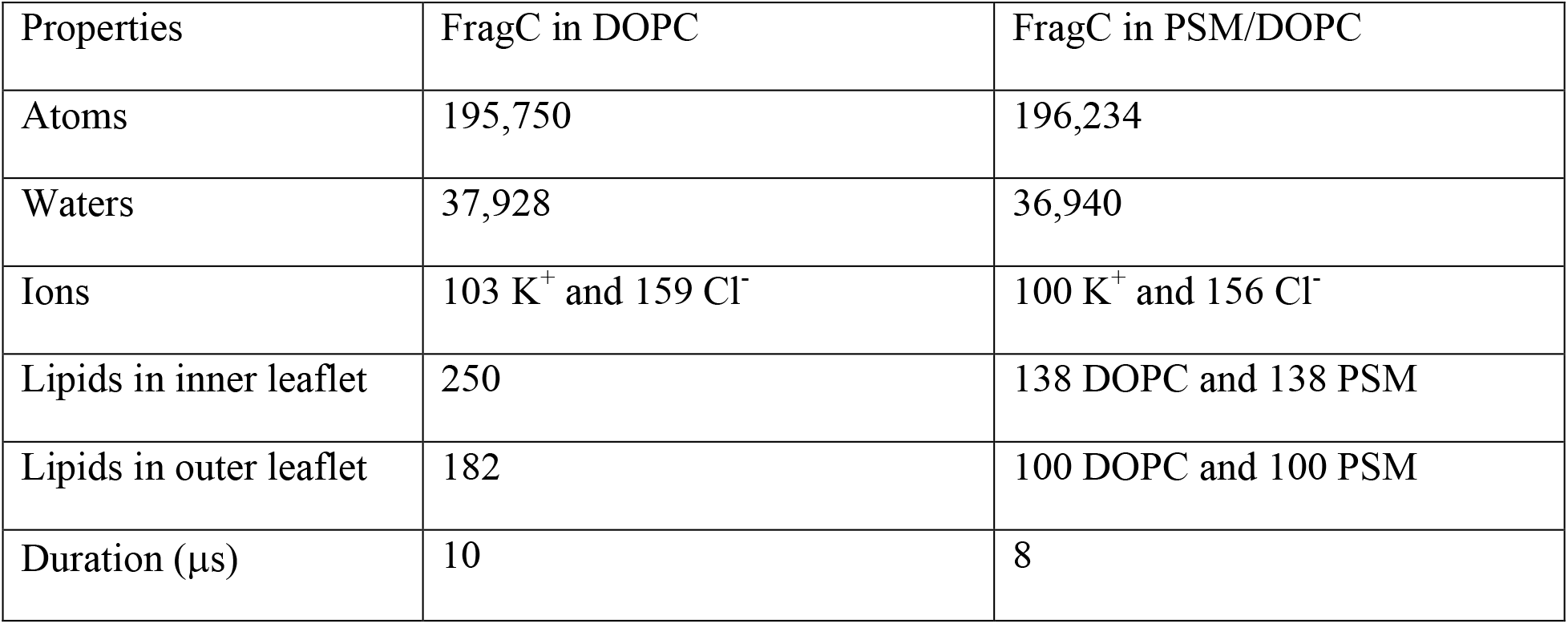
Systems simulated

We calculated pore radii along the *z* axis based on the water volume in the pore using the COOR SEARCH command of the CHARMM program (38) and chose the minimum value as the pore radius (39–41). We use the CHARMM INTE command to calculate short range pairwise interaction energies of peptide-peptide, peptide-lipid and lipid-lipid. Uncertainties were calculated using the method of block averages (42).

For the second system, where two lipid types (i.e. DOPC and PSM) are present, we calculated the following five properties to probe phase coexistence of lipids: a) the average of deuterium order parameters over methylene (-CH_2_-) groups in lipid tails according to this equation:

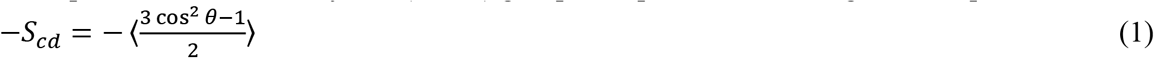

where *θ* is the angle between the *z* axis and the C-H bond. b) Lipid tail tilt angle, defined as the angle between the *z* axis and a vector passing the carbonyl carbon and the terminal carbon of the acyl chain. Both acyl chains in DOPC have carbonyl groups, but only one acyl chain in PSM has a carbonyl group. For the other chain in PSM, we consider the carbon in the C-OH group instead. c) The bilayer thickness calculated as the distance between lipid pairs in two leaflets. Each lipid pair is selected in a way that their head to head distance is minimum. d) The time average of lipid density calculated for each lipid type on each leaflet according to the positions of headgroups. e) The mean square displacement for each lipid type on each leaflet according to movements of headgroups. We used frames of the trajectory spaced at 10 ns to calculate these five properties.

## RESULTS

The octameric complex maintains its structure very well during the 8 or 10 μs of simulation in the two bilayers. The backbone root mean square deviation (RMSD) of the entire complex is 2.7 ± 0.1 Å and 2.6 ± 0.2 Å in DOPC and PSM/DOPC, respectively (Fig. S1a). The RMSD of individual subunits is 2.2 ± 0.4 Å and 2.2 ± 0.3 Å in DOPC and PSM/DOPC, respectively (Fig. S1b). These values were calculated over the last two microseconds of each simulation. The first three missing residues were initially built in an extended conformation (Fig. 2, left column) but most became helical during the simulations. Three N-termini in DOPC (Fig. 2, top right) and one in PSM/DOPC (Fig. 2, bottom right) remained unfolded.

**Fig. 2.**
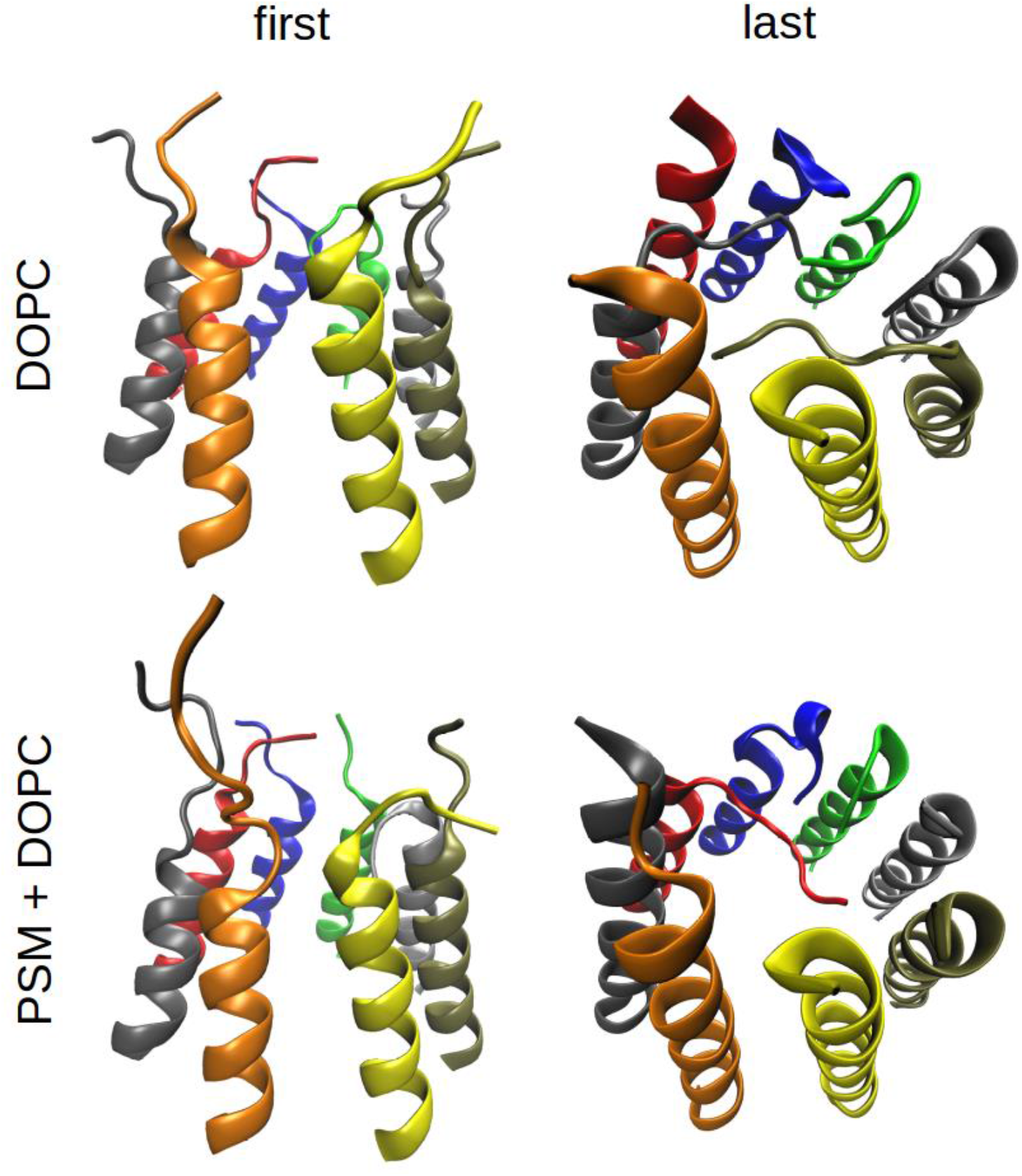
The first 20 residues of each protomer in FragC octamer, in DOPC at the beginning of the simulation (top left), in PSM/DOPC at the beginning of the simulation (bottom left), in DOPC after 10 μs (top right), and in PSM/DOPC after 8 μs (bottom right).

In the initial structure, the pore has a radius of ~7.5 Å, consistent with the 8 Å reported by Tanaka et. al. (7). For both systems, the pore radius decreases slightly over time and fluctuates significantly, due to movements of the unfolded N-termini (Fig. 3). Since there are more unfolded N-termini in DOPC than in PSM/DOPC, fluctuations are higher in DOPC. Over the last 2 μs of each simulation, the pore radius is 4.9 ± 0.2 Å and 5.5 ± 0.05 Å in DOPC and PSM/DOPC, respectively. The number of water molecules inside the FragC pore over the last 2 μs of each simulation is 425 ± 1 and 515 ± 4 in DOPC and DOPC/PSM, respectively.

**Fig. 3.**
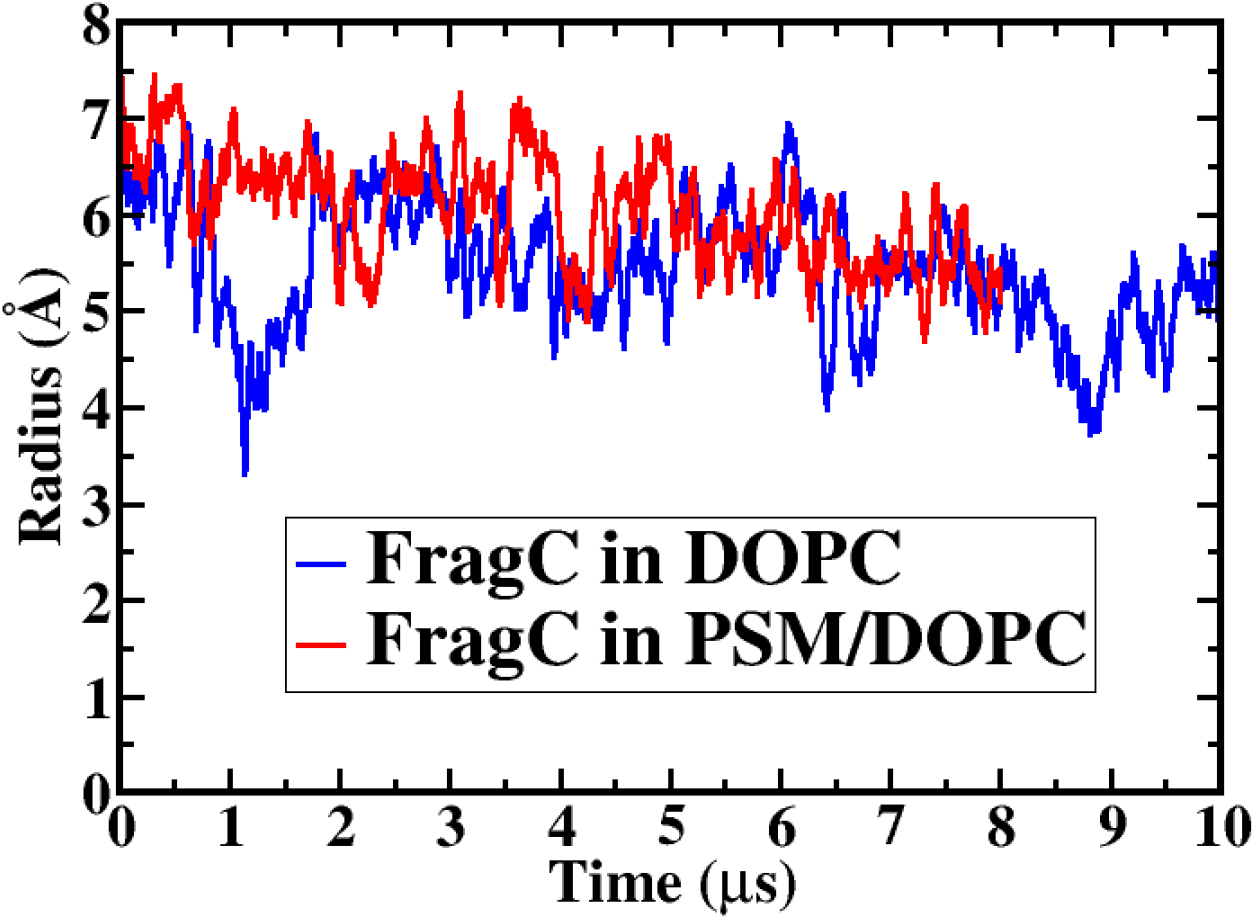
Pore radius of FragC octamer in DOPC and PSM/DOPC lipid bilayers as a function of time

The FragC octamer in both systems forms a funnel-like pore in which the pore size is maximal at the outer leaflet (due to the bulky extramembranous domains) and minimal at the inner leaflet. As a result, there are openings (fenestrations) between adjacent peptides close to the outer leaflet where the α-helix (residues 1-29) is connected to the β-sheet rich region (residues 30-179). These fenestrations are covered by lipids. In the first system, the lipid is necessarily DOPC, but it may be DOPC or PSM in the second system (Fig. S2). Table 2 lists the type of the lipid between each two adjacent peptides in the PSM/DOPC bilayer, the approximate time in the simulation when the lipid headgroup was localized between the peptide pair and the interaction energy of each lipid with the associated peptide pair over the last microsecond of the simulation. When a lipid headgroup enters a fenestration, it is very unlikely to leave it. During the simulation, we have only once observed a DOPC headgroup leave the fenestration between peptides H-A and it was almost immediately substituted by another DOPC headgroup at 0.8 μs. Before the insertion of a lipid headgroup, the fenestration is covered by lipid tails. Throughout the simulation, the fenestration between F and G peptides was covered by lipid tails rather than a headgroup.

**Table 2.**
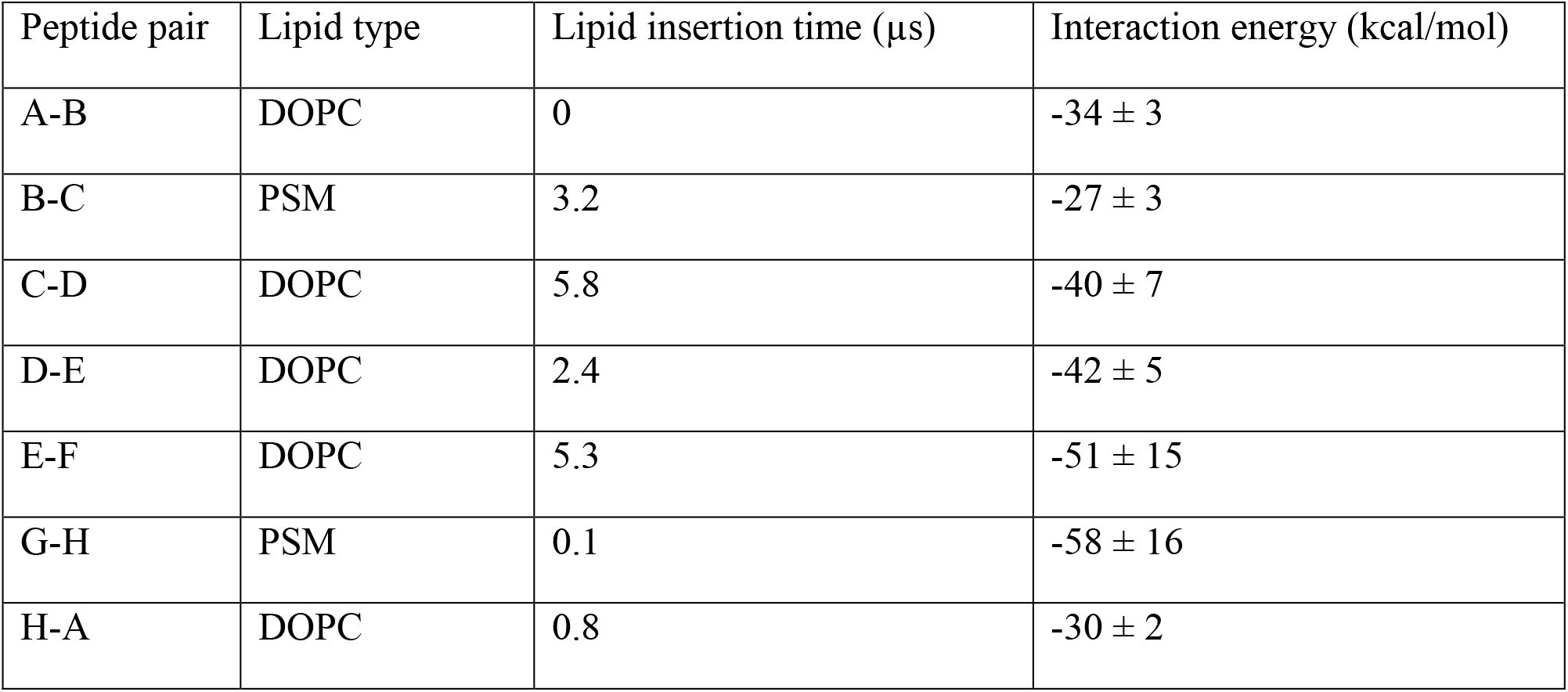
Lipids at each fenestration for FragC in PSM/DOPC and their interaction energy with the two peptides that form the fenestration.

Tanaka et. al. (7) suggested that the lipid inside each fenestration should be a SM because of the strong interactions between the lipid and residue Arg31. However, our results in Table 2 show that there is no preference for PSM in the fenestrations and interaction energies between each peptide pair and their associated lipid do not demonstrate any advantage for PSM. It is true that the strongest interaction energy is observed for a PSM headgroup at G-H, but the difference from the strongest DOPC lipid is smaller than the statistical uncertainty. Furthermore, the groups that differ between the two lipid types are not engaged in h bonding interactions with the peptides. We have also calculated interaction energies between residue Arg31 of each peptide and DOPC or PSM lipids in the outer leaflet over the last microsecond of the simulation (Table 3). Again, the largest interaction energy observed is with PSM (for peptide G), but interactions with DOPC predominate because DOPC concentration around the protein is higher in the outer leaflet (see below).

**Table 3.** Interaction energies (kcal/mol) between Arg31 of each peptide and DOPC or PSM over the last microsecond of the simulation.

| Peptide | DOPC | PSM |
| --- | --- | --- |
| A | $-13.3 \pm 1.5$ | $0.08 \pm 0.02$ |
| B | $7.4 \pm 1.7$ | $-3.4 \pm 1.4$ |
| C | $-21.2 \pm 4.7$ | $0.08 \pm 0.04$ |
| D | $-18.7 \pm 3.5$ | $0.03 \pm 0.01$ |
| E | $-29.2 \pm 0.5$ | $-3.3 \pm 0.2$ |
| F | $-2.4 \pm 0.3$ | $0.06 \pm 0.18$ |
| G | $0.04 \pm 0.01$ | $-33.9 \pm 2.5$ |
| H | $-5.2 \pm 1.7$ | $-2.8 \pm 0.06$ |

An important role of some aromatic residues in the β7-β8 loop (e.g. W112, Y113, etc.) in binding sphingomyelin has been suggested experimentally (18). We have calculated the interaction energies for some of these aromatic residues with DOPC and PSM in the outer leaflet over the first and the last microseconds of the simulation (Table 4). For most of these residues, their interactions with PSM become weaker over time and their interactions with DOPC become stronger. This suggests that PSM gradually moves away from the protein and is replaced by DOPC.

**Table 4.** Interaction energies (kcal/mol) between selected aromatic residues with PSM (E^PSM^) and DOPC (E^DOPC^) during the first and the last microsecond of the simulation.

| Residue | First microsecond |  | Last microsecond |  |
| --- | --- | --- | --- | --- |
| | $E^{\text{PSM}}$ | $E^{\text{DOPC}}$ | $E^{\text{PSM}}$ | $E^{\text{DOPC}}$ |
| W112 | $-78 \pm 3$ | $-83 \pm 4$ | $-68 \pm 5$ | $-114 \pm 5$ |
| W116 | $-45 \pm 3$ | $-31 \pm 4$ | $-20 \pm 3$ | $-42 \pm 3$ |
| Y108 | $-34 \pm 3$ | $-85 \pm 8$ | $-64 \pm 3$ | $-109 \pm 3$ |
| Y113 | $-99 \pm 3$ | $-95 \pm 2$ | $-80 \pm 5$ | $-123 \pm 7$ |
| Y133 | $-25 \pm 3$ | $-23.7 \pm 0.4$ | $-22 \pm 2$ | $-20 \pm 3$ |
| Y137 | $-86 \pm 3$ | $-71 \pm 5$ | $-54 \pm 8$ | $-105 \pm 7$ |
| Y138 | $-67 \pm 3$ | $-52 \pm 3$ | $-55 \pm 5$ | $-64 \pm 3$ |

Fig. 4a shows probability distributions of -S_cd_ for DOPC and PSM lipids in the inner leaflet in the PSM/DOPC simulation over time. Each plot represents the average order parameter distribution of CH_2_ groups of lipid tails over 1 μs (for instance “DOPC 4 μs” means calculated over DOPC lipids between 3 μs and 4 μs). The distribution of DOPC’s order parameter remains unchanged over the simulation whereas PSM’s order parameter distribution moves towards higher values. In other words, PSM lipid tails become more ordered as the simulation progresses. On the other side, the probability distributions of -S_cd_ for DOPC and PSM lipids in the outer leaflet do not show significant change over the simulation time (Fig. 4b).

**Fig. 4.**
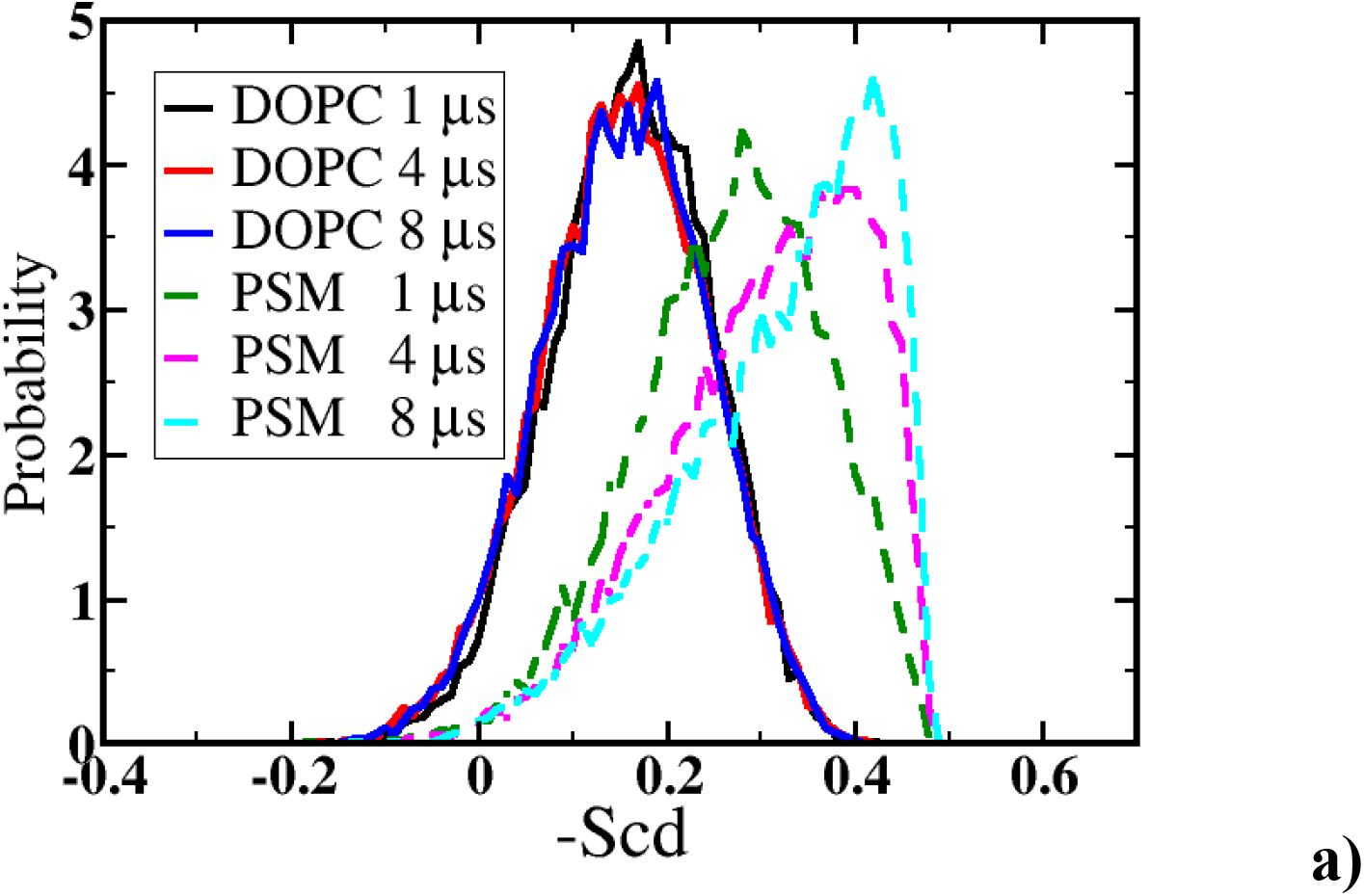
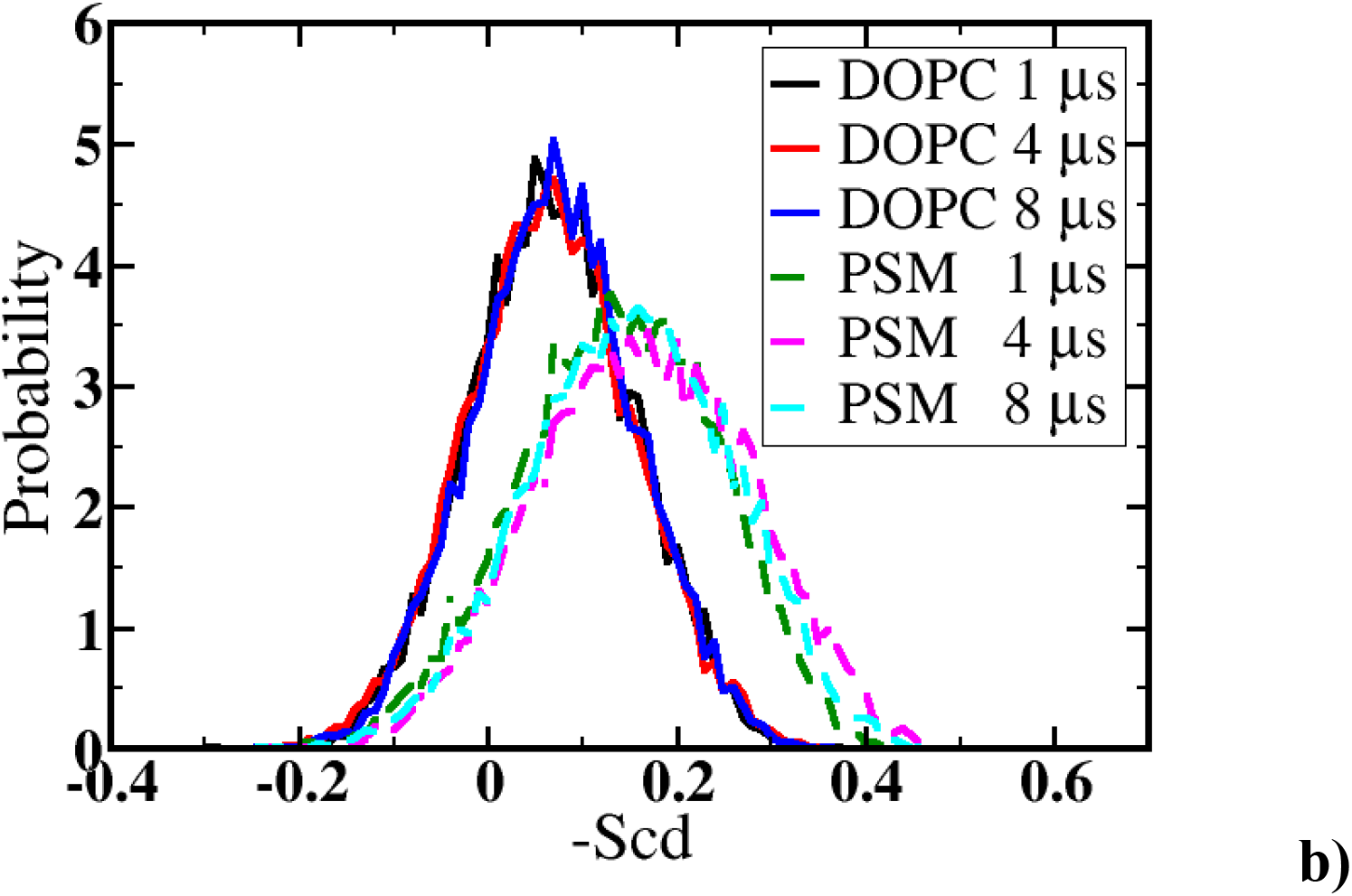
Order parameter distributions of PSM and DOPC at different times: a) inner leaflet, b) outer leaflet

The distributions of lipid tail tilt angles for DOPC and PSM in the inner leaflet (Fig. 5a) indicate that PSM lipids become more aligned with the bilayer normal over time whereas DOPC remains unchanged. On the other side, lipid tail tilt angle distributions in the outer leaflet are unchanged over the simulation for both lipid types (Fig. 5b).

**Fig. 5.**
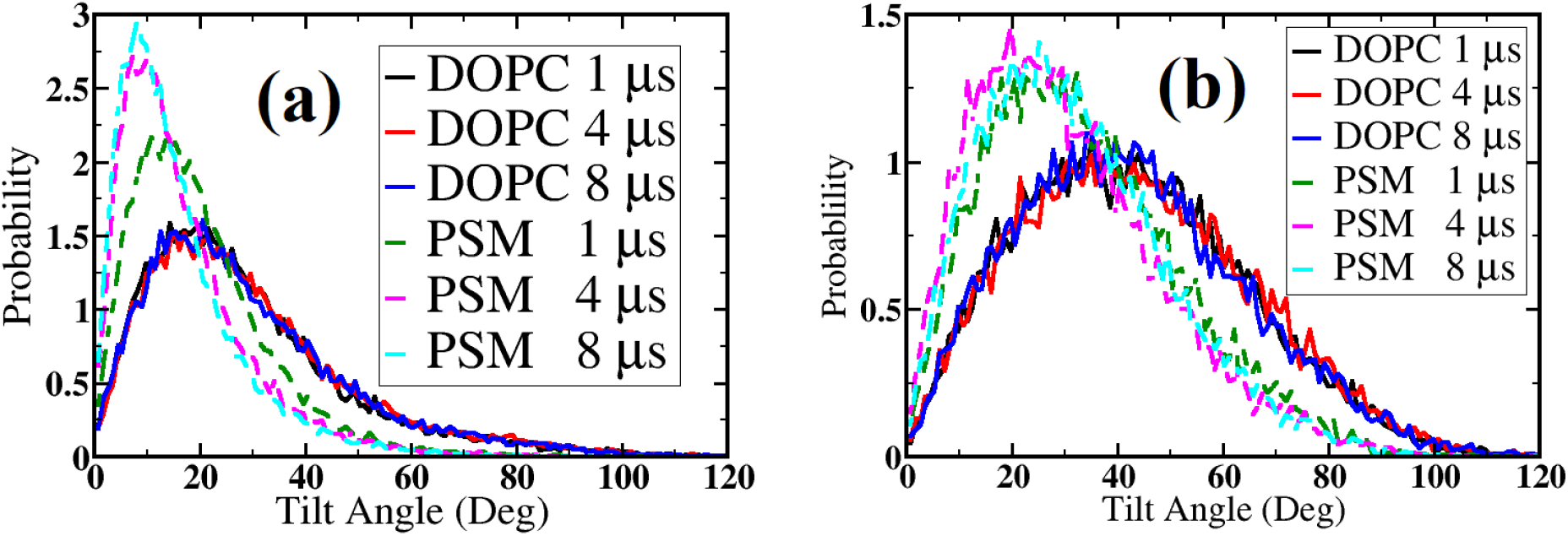
Lipid tail tilt angle distributions of PSM and DOPC at different times. (a) inner leaflet (b) outer leaflet.

Mean square displacements (MSD) of DOPC and PSM headgroups in the inner and outer leaflets are shown in Fig. S3. It seems that the average lipid displacement on each leaflet is independent of lipid type. It is also inferred that lipid displacement is slightly greater in the outer leaflet than in the inner leaflet. Fig. S4 shows lipid bilayer thickness distributions for FragC octamer in PSM/DOPC for the first and the last microsecond of the simulation. We have calculated these values according to lipids in the outer leaflet because there are fewer lipids on this side (200 vs 276 in the inner leaflet). Each panel includes 20,000 points that are all data points we have calculated over one microsecond (i.e. 200 lipids × 100 snapshots). It can be seen that the lipid bilayer is thinner around the pore and thicker away from the pore at corners of the simulation box.

Figs. 6 and 7 show lipid densities for DOPC and PSM over time in the inner and outer leaflets, respectively. The color bars of these two figures show that lipid density reaches higher values in the outer leaflet because the β7-β8 loop of FragC lowers the available area for lipids. So, lipids become more compact. Lipid densities for DOPC and PSM are quite uniform in the beginning of the simulation and become less so over the course of the simulation because lipids of the same type tend to congregate. This is clearer in the inner leaflet (Fig. 6 for the first and the last microsecond of the simulation), where near the end of the simulation PSM tends to concentrate close to the protein and DOPC away from it. The opposite is observed in the outer leaflet (Fig. 7 for the first and the last microsecond of the simulation).

**Fig. 6.**
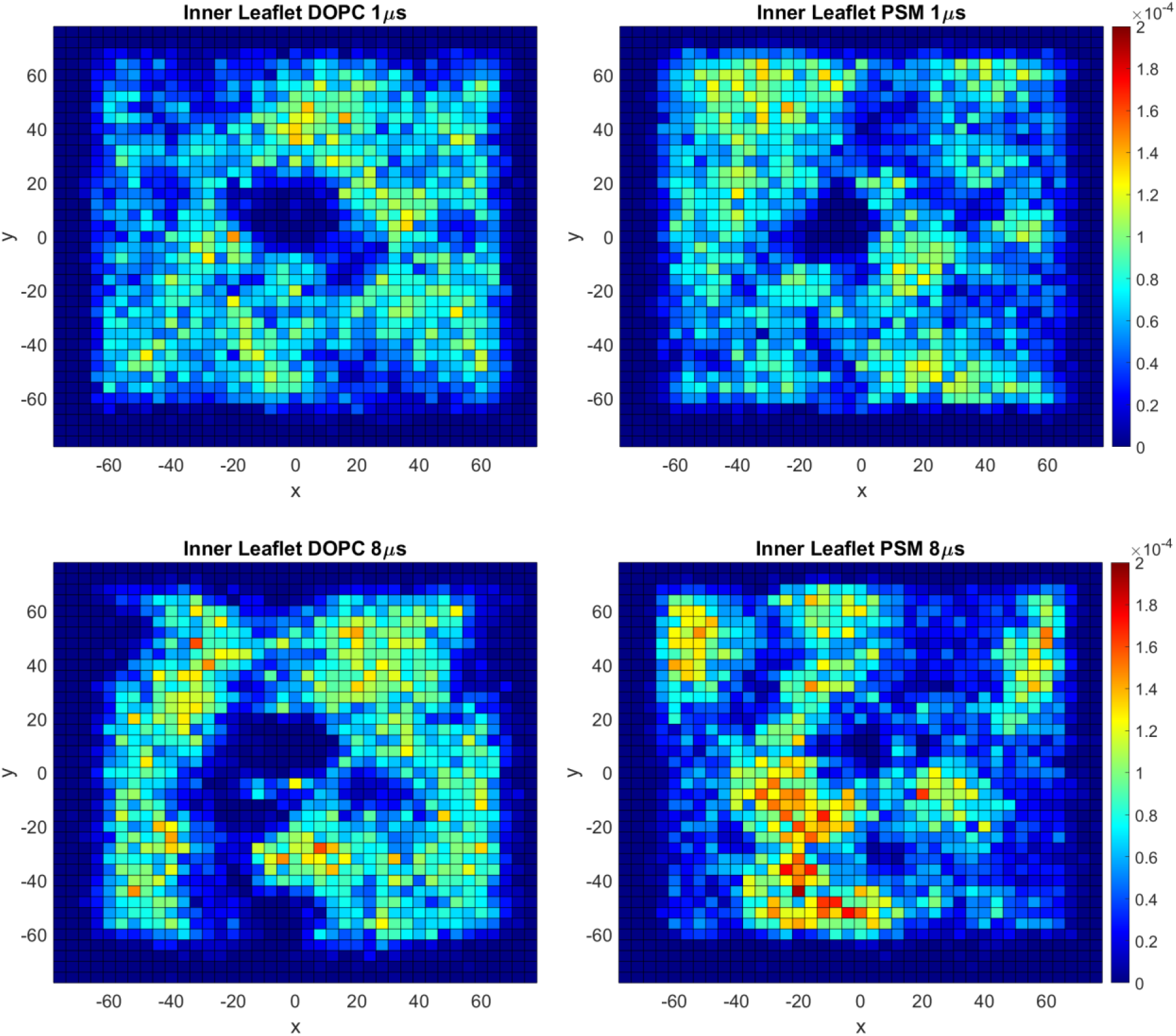
Lipid densities of DOPC and PSM in the inner leaflet for the first and the last microsecond of the simulation. The grid size is 4 Å.

**Fig. 7.**
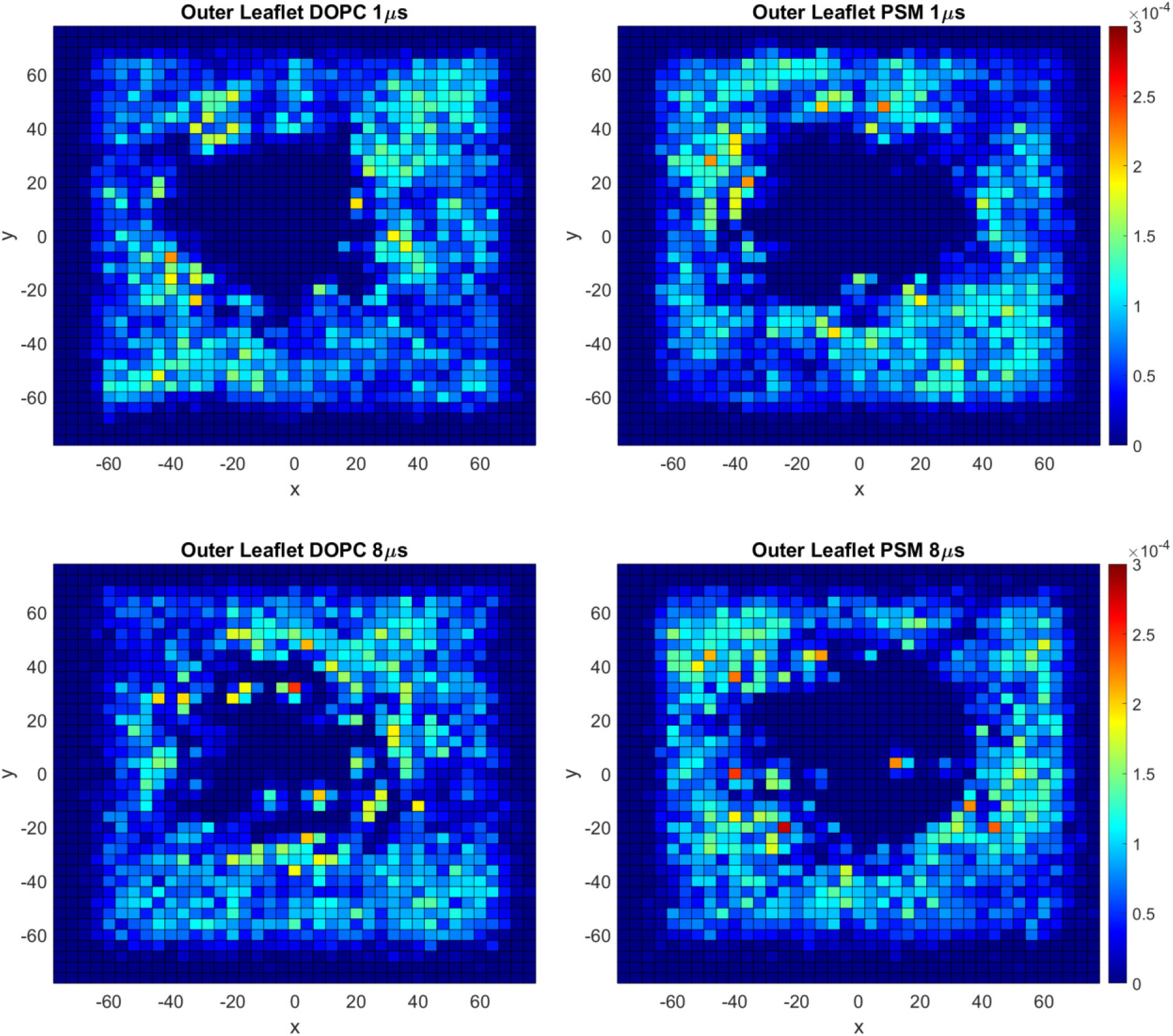
Lipid densities of DOPC and PSM in the outer leaflet for the first and the last microsecond of the simulation. The grid size is 4 Å.

This is corroborated by calculation of interaction energies and radial distribution functions (RDFs) in the PSM/DOPC system. The interaction energies of Ser1 with lipid headgroups are −81 ± 6 kcal/mol and −48 ± 10 kcal/mol for PSM and DOPC, respectively. RDF plots of Ser1 and PSM or DOPC (Fig. 8) also show that over the simulation, more PSM lipids surround Ser1 than DOPC. In contrast, in the outer leaflet the protein is surrounded mostly by DOPC. Fig. 9 shows RDF plots of three aromatic residues from Table 4 with PSM and DOPC during the first and last microseconds of the simulation (Fig. 9). Table 4 and Fig. 9 show that in the outer leaflet FragC becomes more surrounded by DOPC as the simulation progresses.

**Fig. 8.**
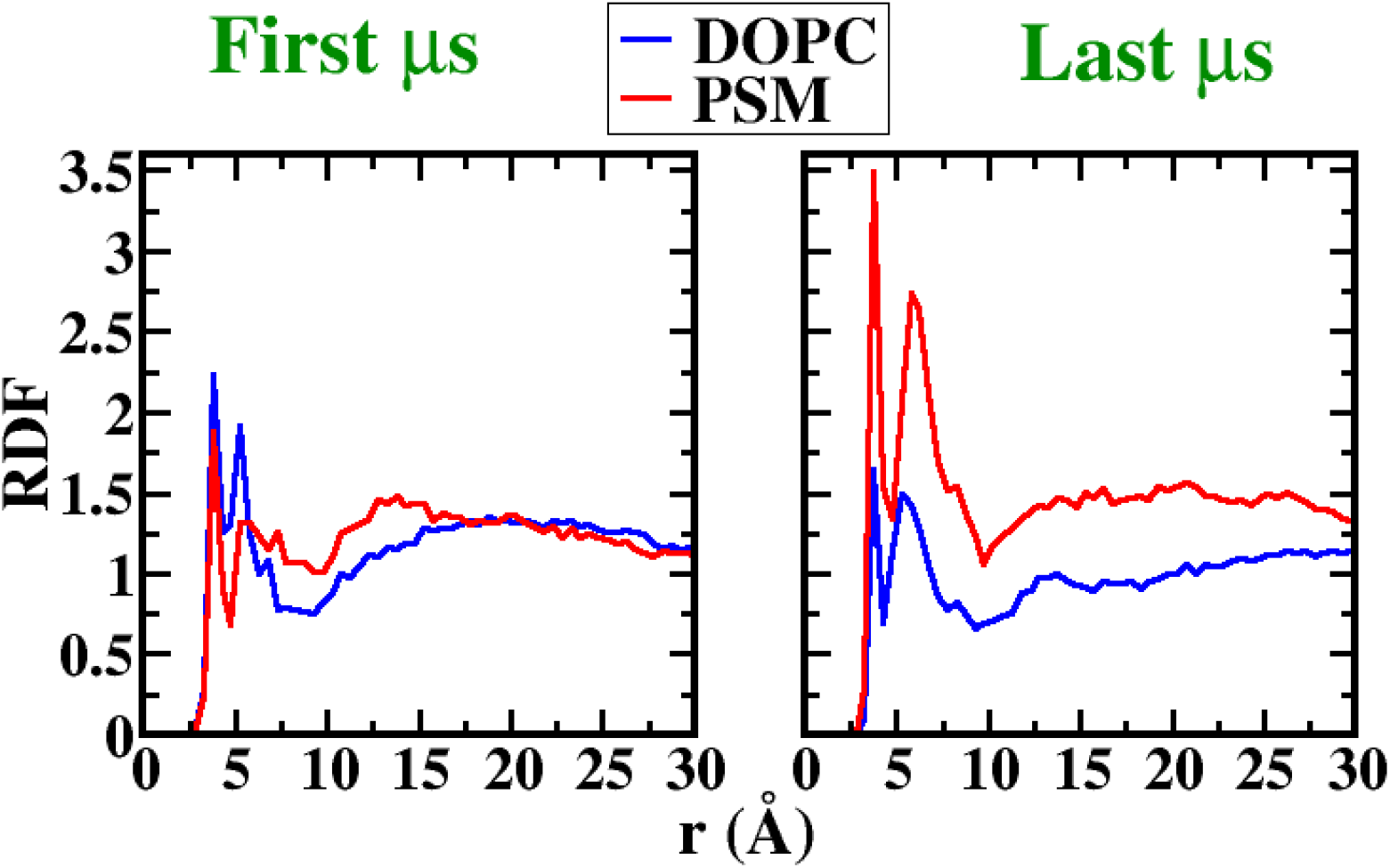
RDF plots of Ser1 and lipid headgroups (PSM/DOPC) during the first and last microseconds of the simulation.

**Fig. 9.**
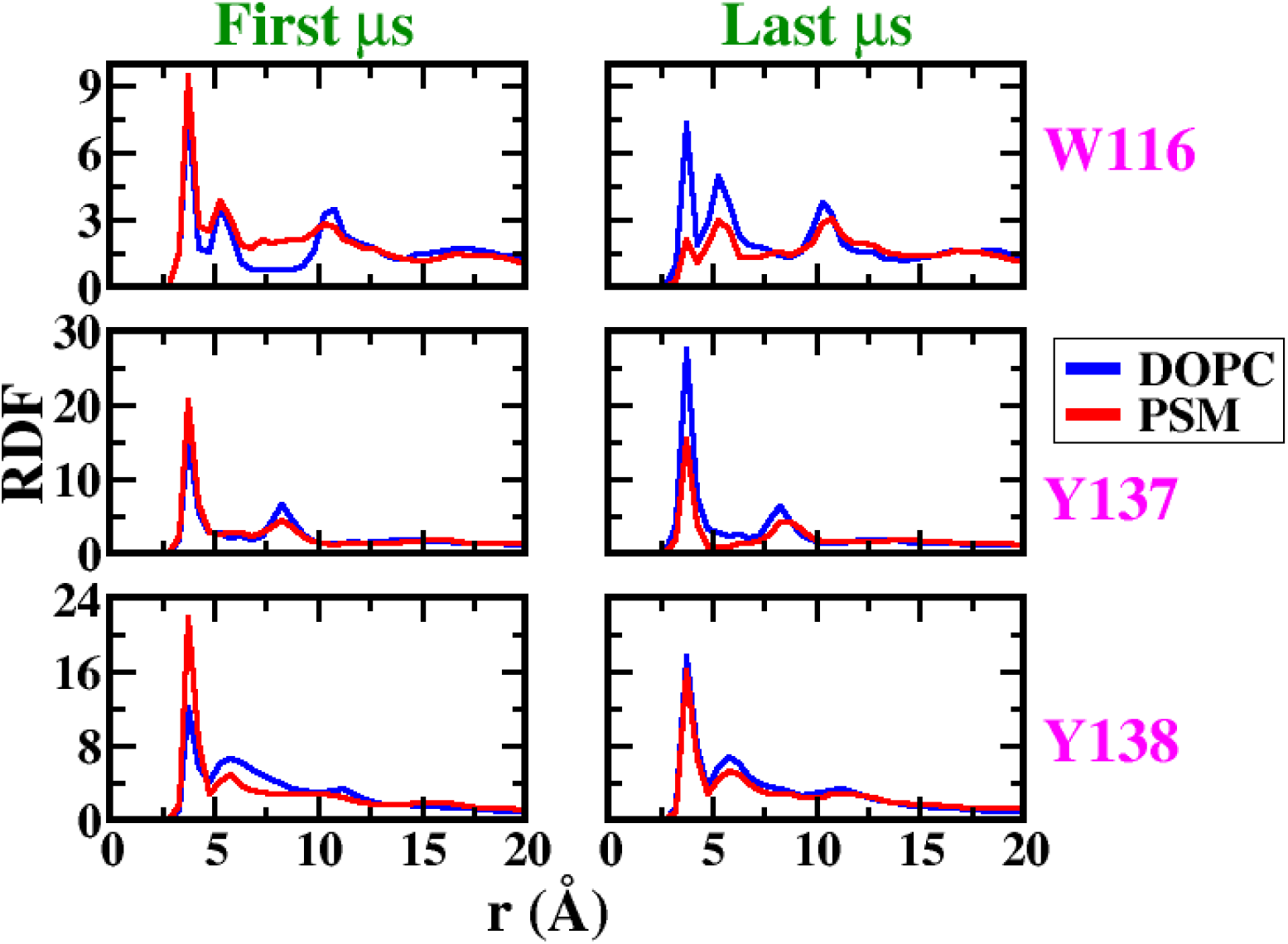
RDF plots of selected aromatic residues and lipid headgroups (PSM/DOPC) during the first and last microseconds of the simulation.

Table 5 lists protein-lipid and lipid-lipid interaction energies over the first and last microseconds of the simulation in both leaflets. Protein-lipid interactions are weaker in the inner leaflet because they involve only one half of the N-terminal helices. During the simulation, there is a slight increase in protein-PSM interactions in the inner leaflet and a substantial decrease in interactions with DOPC. The opposite is observed in the outer leaflet. Lipid-lipid interactions change dramatically in the inner leaflet, with DOPC-PSM interactions decreasing and PSM-PSM and DOPC-DOPC interactions increasing. This points to a tendency for phase separation in the inner leaflet. In the outer leaflet, lipid-lipid interactions change more subtly. Cross-interactions decrease slightly and PSM-PSM interactions increase slightly, but DOPC-DOPC interactions also tend to decrease.

**Table 5.**
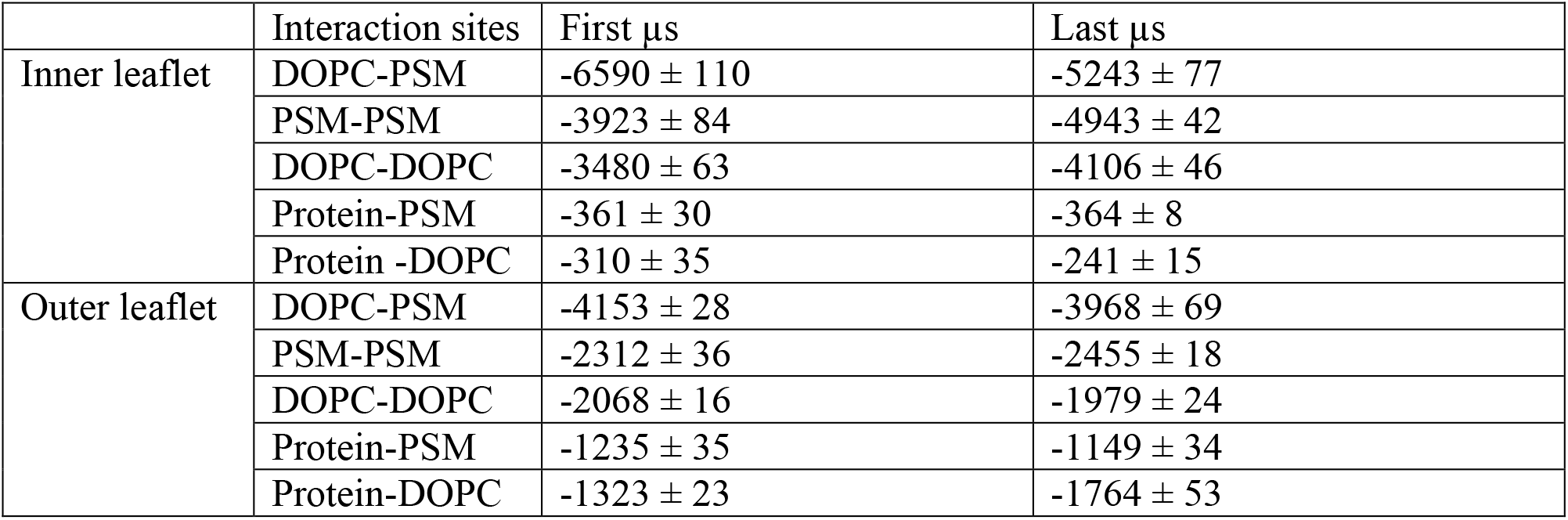
Lipid-lipid and protein-lipid interaction energies (kcal/mol) in the inner and outer leaflets.

| | Interaction sites | First μs | Last μs |
| --- | --- | --- | --- |
| Inner leaflet | DOPC-PSM | $-6590 \pm 110$ | $-5243 \pm 77$ |
| | PSM-PSM | $-3923 \pm 84$ | $-4943 \pm 42$ |
| | DOPC-DOPC | $-3480 \pm 63$ | $-4106 \pm 46$ |
| | Protein-PSM | $-361 \pm 30$ | $-364 \pm 8$ |
| | Protein-DOPC | $-310 \pm 35$ | $-241 \pm 15$ |
| Outer leaflet | DOPC-PSM | $-4153 \pm 28$ | $-3968 \pm 69$ |
| | PSM-PSM | $-2312 \pm 36$ | $-2455 \pm 18$ |
| | DOPC-DOPC | $-2068 \pm 16$ | $-1979 \pm 24$ |
| | Protein-PSM | $-1235 \pm 35$ | $-1149 \pm 34$ |
| | Protein-DOPC | $-1323 \pm 23$ | $-1764 \pm 53$ |

## DISCUSSION

The present simulations showed that the FragC octamer and its pore structure are very stable in both DOPC and PSM/DOPC lipid bilayers on a multi-microsecond time scale. More helix fraying was observed in DOPC, but this may be incidental. The pore radius is essentially indistinguishable in the two bilayers, although the pore contains somewhat more water molecules in the mixed bilayer. We observe different behavior of the lipids in the two bilayer leaflets. The protein tends to be surrounded by PSM in the inner leaflet and DOPC in the outer leaflet. We found no meaningful preference for a lipid type in the fenestrations where the N-terminal α-helix connects to the extramembranous domains.

Strong homophilic interactions were observed between lipids, especially between PSM, that seem to drive a phase transition for PSM in the inner leaflet but not in the outer leaflet. This may be because protein-lipid interactions in the outer leaflet prevent or slow down phase separation. The fact that there is a greater number of lipids in the inner leaflet may also play a role. Experimentally, an equimolar DOPC/PSM bilayer exhibits coexistence of a liquid disordered and a gel phase between 8 and 37 ^o^C (43), while pure PSM exhibits a transition to the liquid phase at 41 ^o^C (43, 44). Our simulations are at 37 ^o^C, thus it is not unreasonable to observe phase separation. To our knowledge, the melting temperature of PSM in the force field used here has not been determined.

One key question in the actinoporin field is membrane targeting and the role of sphingomyelin. One school of thought is that SM is recognized specifically by aromatic residues in membrane-binding loops of the core domain (2). This is supported by the fact that preincubation with SM inhibits the toxin (16), measurements showing that it binds only to SM-containing vesicles (18), and the effect of mutations of aromatic residues on toxin action (18). It is hard to imagine how this recognition of SM over PC takes place from above the membrane, since they both expose the same headgroup. For another SM-dependent toxin, lysenin, a crystal structure showed both the headgroup and acyl chain of an individual SM molecule bound to the protein (45). However, this SM molecule had to be extracted from a lipid bilayer to be able to bind in the designated way. An alternative theory is that SM acts indirectly, via inducing phase coexistence (20, 22). This is supported by experiments that show ion conduction (19) and dye leakage (20, 46, 47) in PC/cholesterol mixtures. In pure SM dye leakage by equinatoxin II is reduced compared to an equimolar PC/SM mixture (20, 47).

The present simulations support the latter view. We find that in the outer leaflet the protein interacts preferentially with DOPC, not PSM. This seems consistent with the finding that the final complex resides in the liquid disordered phase, where DOPC would be dominant (23). Our results, however, are silent on the issue of why phase coexistence promotes actinoporin function. The present work examines the final pore state. Phase coexistence may affect earlier steps, such as membrane binding, oligomerization, conformational change, and Nt helix insertion.

## AUTHOR CONTRIBUTIONS

AS and BN performed the simulations. AS analyzed the simulations. AS and TL wrote the manuscript.

## ACKNOWLEDGMENTS

This work was supported by NIH (GM117146) and infrastructure support was provided in part by Research Centers in Minority Institutions grant no. 8G12MD007603 from the NIH. Anton computer time was provided by the Pittsburgh Supercomputing Center through grant R01GM116961 from the NIH. The Anton machine at PSC was generously made available by D.E. Shaw Research, and computer time was provided by the National Center for Multiscale Modeling of Biological Systems through grant no. P41GM103712-S1 from the NIH and Pittsburgh Supercomputing Center.

